# Differential expression of COVID-19-related genes in European Americans and African Americans

**DOI:** 10.1101/2020.06.09.143271

**Authors:** Urminder Singh, Eve Syrkin Wurtele

## Abstract

The Coronavirus disease 2019 (COVID-19) pandemic has affected African American populations disproportionately in regards to both morbidity and mortality. A multitude of factors likely account for this discrepancy. Gene expression represents the interaction of genetics and environment. To elucidate whether levels of expression of genes implicated in COVID-19 vary in African Americans as compared to European Americans, we re-mine The Cancer Genome Atlas (TCGA) and Genotype-Tissue Expression (GTEx) RNA-Seq data. Multiple genes integral to infection, inflammation and immunity are differentially regulated across the two populations. Most notably, F8A2 and F8A3, which encode the HAP40 protein that mediates early endosome movement in Huntington’s Disease, are more highly expressed by up to 24-fold in African Americans. Such differences in gene expression can establish prognostic signatures and have critical implications for precision treatment of diseases such as COVID-19. We advocate routine inclusion of information such as postal code, education level, and profession (as a proxies for socioeconomic condition) and race in the metadata about each individual sampled for sequencing studies. This relatively simple change would enable large-scale data-driven approaches to dissect relationships among race, socio-economic factors, and disease.

## Introduction

As of June 1, 2020, the COVID-19 pandemic has infected over 6.3 million people and killed over 370,000 worldwide (https://coronavirus.jhu.edu/map.html). Its causative agent, the novel SARS-CoV-2, is an enveloped single stranded RNA virus that infects tissues including lung alveoli^1, 2^, renal tubules^3, 4^, the central nervous system^5^, ileum, colon and tonsils^6–8^, and myocardium^9, 10^.

The complex combinations of symptoms and disease caused by SARS-CoV-2 include fever, cough, fatigue, dyspnea, diarrhea, stroke, acute respiratory failure, renal failure, cardiac failure, potentially leading to death^9, 11–14^. Symptoms are induced by direct cellular infection and proinflammatory repercussions from infection in other regions of the body^13–15^. A body of evidence indicates that long-term health implications can follow SARS-CoV-2 infection^16, 17^. The attributes of the human host that impact COVID-19 morbidity and mortality are not well understood^6, 13, 14, 18–23^.

Risk factors for complications of COVID-19 include 65+ years, obesity, and comorbidities such as diabetes, hypertension and heart disease^24^. Heritable factors in the human host influence COVID-19 symptoms^25^. However, to date, only a few of the genetic determinants of COVID-19 severity have been even partially elucidated. Genetic variants of Angiotensin-Converting Enzyme2 (ACE2), the major human host receptor for the SARS-CoV-2 spike protein, may be linked to increased infection by COVID-19^26–29^. Human Leukocyte Antigen (HLA) gene alleles have been associated with susceptibility to diabetes and SARS-CoV-2^30^. The genetic propensity in southern European populations for mutations in the pyrin-encoding Mediterranean Fever gene (MEFV) may elevate levels of pro-inflammatory molecules, leading to a cytokine storm^31^ and greater severity of COVID-19^32–35^. Identifying those individuals most at-risk for severe COVID-19 infection, and determining the molecular and physiological basis for this risk, would enable more informed public health decisions and interventions.

COVID-19 cases and deaths are disproportionately higher among African Americans. One cause of this disparity are complex socio-economic factors^15, 36, 37^. A number of studies have shown differences in genes expression among races^38–41^. Gene expression reflects the interaction between environmental, physiological, and genetic influences. This study specifically investigates whether the expression of genes that are implicated in the severity of COVID-19 infection varies with race. Understanding differences in expression at the population level could help predict risk factors and identify more personalized, treatments for COVID-19.

Here, we re-mine existing RNA-Seq data and reveal significant differences in expression between European Americans and African American of multiple genes potentially involved in COVID-19-associated inflammation and immunomodulatory reponse.

## Results

We evaluated differential expression by re-mining an aggregated dataset of 7,142 RNA-Seq samples^42^ modified from the normalized and batch-corrected data from the GTEx and TCGA projects^43^. The Genotype-Tissue Expression (GTEx) project provides data representing “non-diseased” conditions from diverse tissues. The well-curated TCGA project is the largest project available with easily accessible metadata on the races of the individuals who contributed samples representing diseased tissues (tumors) of multiple origins. These large data provide a unique opportunity to evaluate differences in gene expression across populations in multiple tissues in diseased and normal states.

Re-mining existing RNA-Seq data and metadata has several caveats. Because race assignments are self-reported, many of the individuals sampled will be from admixed populations^41, 44^. We are terming those self-reporting as “Black or African American” as “African Americans” and “White” as “European Americans”. The ancestry of the preponderance of African Americans is Western Africa^44^, thus our results for African Americans would mostly reflect more specifically Western Africans. Those self-reporting as White are presumed to be predominantly European Americans, but this group also would include individuals of other populations, including Indians and admixed Hispanic individuals, depending on how these individuals chose to self-report.

Also, we were limited to comparison of differences between gene expression in African American and European American populations, because even in these large studies, the sample numbers for the other three major population groups (Asian, Native American, and Pacific Islanders)^45^ were too low for robust statistical assessment. Even between these two populations, not all conditions (cancers or “non-diseased” tissues) had sufficient samplings of African Americans for robust statistical assessment (Table 1; Supplementary Table S1).

**Table 1.** Number of DE genes in African Americans compared to European Americans in eight cancer types and nine non-diseased tissue types. Criteria for DE, >2-fold difference in expression, Mann-–Whitney U test is significant with BH-corrected p-value < 0.05.

| Project | Condition | #AA | #EA | #Upregulated | #Downregulated |
| --- | --- | --- | --- | --- | --- |
| TCGA | Breast invasive carcinoma (BRCA) | 142 | 674 | 83 | 164 |
| TCGA | Colon adenocarcinoma (COAD) | 54 | 188 | 30 | 21 |
| TCGA | Kidney renal clear cell carcinoma (KIRC) | 46 | 410 | 68 | 94 |
| TCGA | Kidney renal papillary cell carcinoma (KIRP) | 49 | 166 | 19 | 13 |
| TCGA | Lung adenocarcinoma (LUAD) | 48 | 368 | 16 | 5 |
| TCGA | Lung squamous cell carcinoma (LUSC) | 28 | 337 | 2 | 0 |
| TCGA | Thyroid carcinoma (THCA) | 25 | 292 | 3 | 3 |
| TCGA | Uterine Corpus Endometrial Carcinoma (UCEC) | 54 | 70 | 28 | 5 |
| GTEX | Breast | 12 | 75 | 0 | 0 |
| GTEX | Colon | 41 | 292 | 13 | 9 |
| GTEX | Esophagus | 80 | 564 | 19 | 11 |
| GTEX | Liver | 15 | 97 | 0 | 0 |
| GTEX | Lung | 39 | 269 | 45 | 20 |
| GTEX | Prostate | 13 | 89 | 0 | 0 |
| GTEX | Stomach | 29 | 159 | 4 | 6 |
| GTEX | Thyroid | 43 | 267 | 25 | 30 |
| GTEX | Uterus | 13 | 68 | 0 | 0 |

Finally, although the GTEx project analyzes non-diseased tissues, many of the individuals who donated tissues were severely ill or postmortem from varied causes, which would likely effect gene expression in these tissues.

We analyzed the data and metadata with MetaOmGraph (MOG), software that supports interactive exploratory analysis of large, aggregated data^42^. Exploratory data analysis uses statistical and graphical methods to gain insight into data by revealing complex associations, patterns or anomalies within that data at different resolutions^46^.

Differentially expressed (DE) genes among samples from European Americans and African Americans were identified in a tumor-specific or tissue-specific manner using MOG (Mann-Whitney U test). Of the tumor types in the TCGA data, BRCA, COAD, KIRC, KIRP, LUAD, LUSC, THCA, and UCEC had sufficient numbers of samples for DE analysis (Table 1). GTEx normal tissues analyzed were: breast, colon, esophagus, lung, liver, prostate, stomach, thyroid, and uterus (Table 1). We define a gene as DE between two groups if it meets each of the following criteria:

1. Estimated fold-change in expression of 2-fold or more (log fold change, |*logFC*|⩾ 1 where *logFC* is calculated as in limma^47^.)
2. Mann-–Whitney U test is significant between the two groups (BH corrected p-value < 0.05)

Supplementary Tables S2-S25 contains the full list of DE genes between African American and European American populations for each condition. The numbers of DE genes vary depending on the condition the samples were obtained from. Many genes follow a similar DE trend in diseased as in non-diseased tissues, however, in each case, the fold-change difference of expression among the DE genes was larger in the cancers than in the corresponding non-diseased tissues.

We investigate the distribution of the gene expression values with two additional statistical analyses. We performed the two-sample Kolmogorov–Smirnov test (KS test) to assess whether there is a significant difference in distribution of gene expression between African Americans and European Americans. Hartigans’ dip test was used to test whether a given distribution shows bi or multi-modality. Bi- or multi-modal distributions indicate there may be hidden or unknown covariates affecting the gene expression. Within a race, this could imply presence of sub-population structure. These analyses indicate the not only are genes differentially expressed between the two populations, but the overall distribution in expression values often differed between the populations, for an obvious example, GSTM1 expression in KIRC (Figure 4. Also, in many cases, one or both populations have a bimodal distribution, for example, GSTM1 expression in BRCA has a bimdodal distribution in European Americans but not in African Americans (Figure 4).

**Figure 1.**
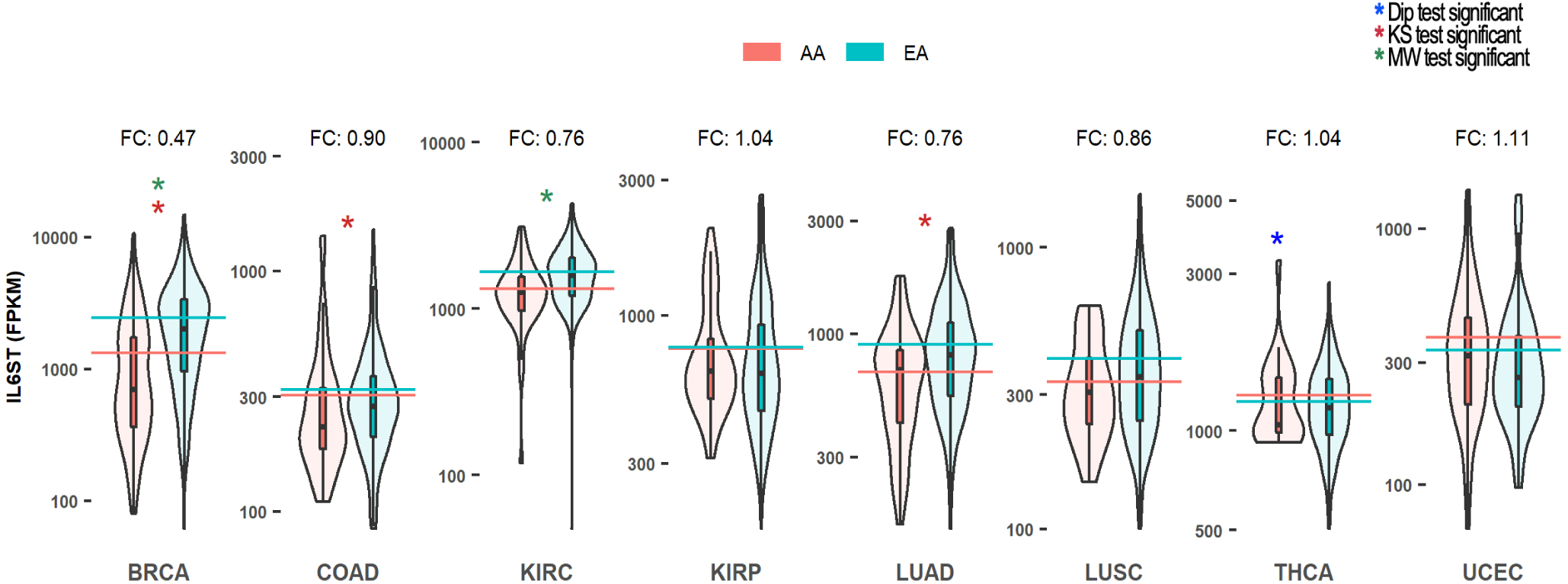
Expression of the **IL6ST** interleukin signal transducer gene in African Americans and European Americans across eight cancer conditions. Violin plots summarizing the expression over each tumor sample in the two populations. AA, African American; EA, European American. Horizontal lines represent mean log expression. *, Hartigans’ dip test significant (p-value < 0.05); *, KS test significant (p-value < 0.05); *, Mann-–Whitney U test significant (BH corrected p-value < 0.05).

**Figure 2.**
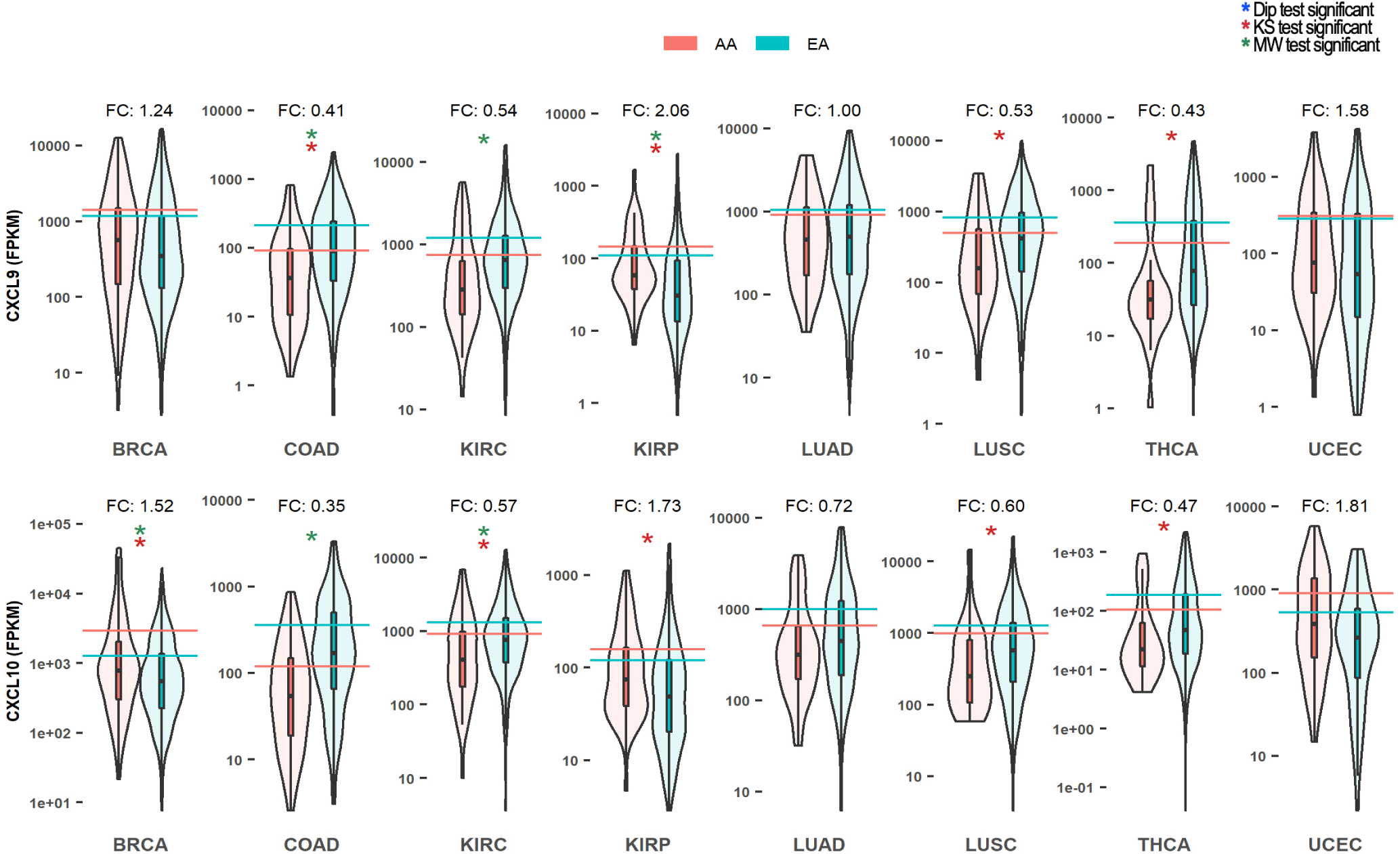
Expression of the **CXCL9** and **CXCL10** circulating chemokine genes in African Americans and European Americans across eight cancer conditions. Violin plots summarize the expression over each tumor sample in the two populations. AA, African American; EA, European American. Horizontal lines represent mean log expression. *, Hartigans’ dip test significant (p-value < 0.05); *, KS test significant (p-value < 0.05); *, Mann-–Whitney U test significant (BH corrected p-value < 0.05).

**Figure 3.**
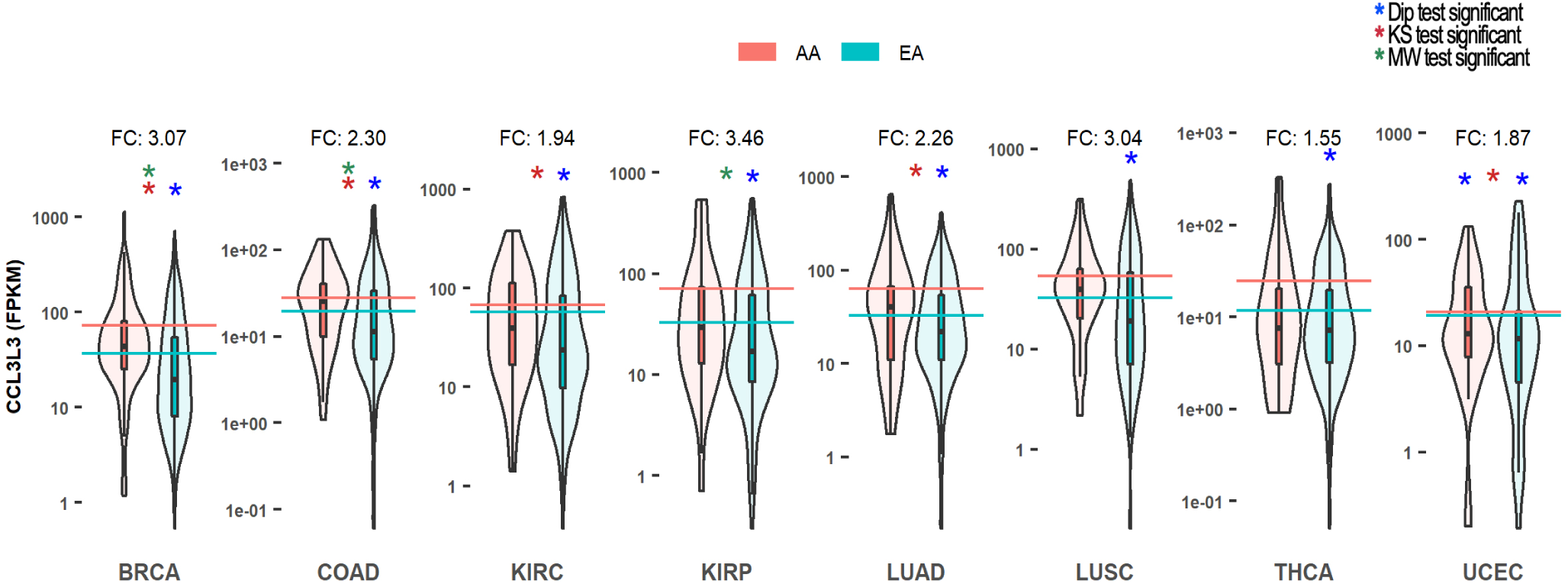
Expression of the **CCL3L3** chemokine gene in African Americans and European Americans across eight cancer conditions. AA, African American; EA, European American. Horizontal lines represent mean log expression. *, Hartigans’ dip test significant (p-value < 0.05); *, KS test significant (p-value < 0.05); *, Mann-–Whitney U test significant (BH corrected p-value < 0.05).

**Figure 4.**
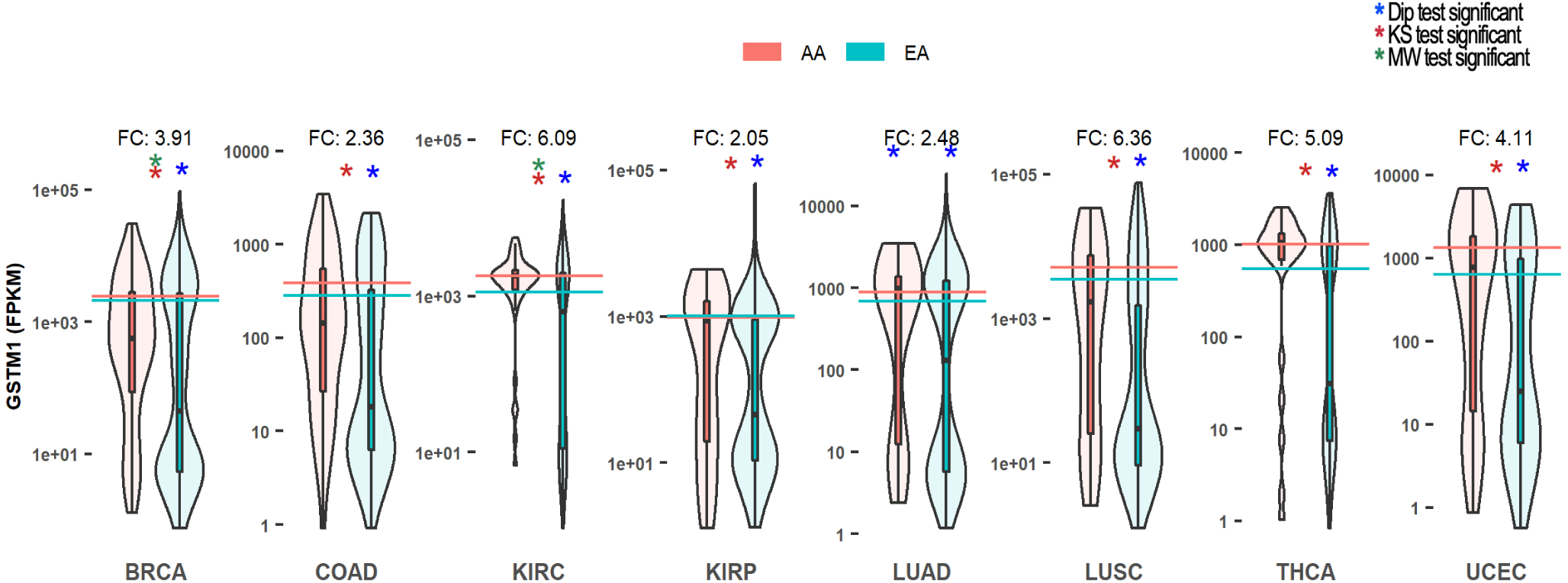
Expression of the mitochondrial glutathione-S-transferase gene, **GSTM1** in African Americans and European Americans across eight cancer conditions. GSTM1 is a key player in metabolism of ROS. Violin plots summarize GSTM1 expression over each tumor sample in the two populations. AA, African American; EA, European American. Horizontal lines represent mean log expression. *, Hartigans’ dip test significant (p-value < 0.05); *, KS test significant (p-value < 0.05); *, Mann-–Whitney U test significant (BH corrected p-value < 0.05).

GO terms related to the biological processes of infection, inflammation and immunity are overrepresented among the genes differentially expressed between African Americans and European Americans. Table 2 lists these GO terms. Supplementary Table S29 provides the overrepresented Gene Ontology (GO) terms among DE genes for each diseased or non-diseased tissue. We drew from *in silico* studies^48^ and experimental analyses especially human responses to infection by SARS-CoV-2 and other coronaviruses^49–52^ to identify ten genes implicated in cellular responses to SARS-CoV-2 infection that are among those differentially expressed in African American and European American populations. Molecular functions of these genes include receptor kinases, cytokines, other signal transduction molecules, and antioxidants. These genes are integral to central cellular processes that affect pathogenesis by SARS-CoV-2, including endosomal development, autophagy, immunity and inflammation^6, 51, 53^. Molecular functions of these genes include receptor kinases, cytokines, other signal transduction molecules, and antioxidants.

**Table 2.** GO terms related to infection, inflammation and immunity that are most enriched among genes that are DE in African Americans compared to European Americans in cancers and non-diseased tissue types.

| Tissue | GO Term | Fold Enrichment | FDR |
| --- | --- | --- | --- |
| COAD | humoral immune response | 17 | 0.003 |
| KIRC | positive regulation of fibrinolysis | 76 | 0.001 |
| LUSC | immune response mediated by microbial peptide | 11 | 0.04 |
| THCA | regulation of T-Cell migration | 23 | 0.03 |
| THCA | killing of cells of other organisms | 14 | 0.03 |
| THCA | antimicrobial humoral response | 10 | 0.03 |
| THCA | receptor-mediated endocytosis | 8 | 0.06 |
| Lung | glutathione derivative biosynthesis) | 71 | 0.001 |
| Esophagus | cellular detoxification of nitrogen) | >100 | 0.04 |

### Cytokines and the storm

A number of genes of the immune response are differentially expressed between African American and European American populations. Expression of IL6ST, a component of the cytokine receptor complex that acts as signal transducer for cytokine interleukins IL6 and IL7, is 2-fold lower in African Americans than European Americans in BRCA (Figure 1).

Circulating chemokines CXCL9 and CXCL10 are also differentially expressed in African Americans as compared to European Americans in several cancers. CXCL9 expression is 2-fold greater in KIRP and over 2-fold lower in COAD and KIRC; CXCL10 expression is 1.5-fold higher in BRCA, and over 2-fold lower in COAD, KIRC and TCHA (Figure 2).

The small inducible chemokine CCL3L3 is upregulated in African Americans by 2-to 3-fold in BRCA, COAD, and KIRP (Figure 3) and in several non-diseased tissues (Supplementary Table S16-S25).

Carcinoembryonic Antigen-related Cell Adhesion Molecules CEACAM5 and CEACAM6 are both downregulated 2 to 3-fold in BRCA in African Americans. (Supplementary Table 2).

### Reactive Oxygen Species

Expression of GSTM1, a key enzyme involved in oxidative stress, differs between African Americans and European Americans. Expression is up to to 6-fold higher in African Americans in the cancers we evaluated (Figure 4). Distribution of expression in European Americans, but not African Americans is bimodal. GSTM1 expression is also DE in non-diseased esophagus and thyroid gland (Supplementary Table S18, S24).

### F8As, endocytosis, and autophagy

Endocytosis and autophagy are intimately interrelated with Covid-19^54^. One little-studied player implicated in early endosome motility^55^ and hence the endocytotic pathway and autophagy, is the seven tetratricopeptide-like repeat F8A/HAP40 (HAP40) protein^56^. In human genomes, three genes, F8A1, F8A2, and F8A3, encode the HAP40 protein^56^. The F8A genes are located on the X chromosome. F8A1 is within intron 22 of the coagulation factor VIII gene, which has a high frequency of mutations^57^; F8A2, and F8A3 are located further upstream.

F8A1, F8A2 and F8A3 are each differentially expressed in African Americans versus European Americans. F8A1 is more highly expressed by about 2-fold in European Americans in every cancer analyzed (Figure 5) and by 2-fold in non-tumor colon (Supplementary Table S17). Conversely, F8A2 and F8A3 are more highly expressed in African Americans in all cancer types. Expression of F8A2 in African Americans is up to 24-fold greater; expression of F8A3 is up to 6.6-fold greater. In LUSC, F8A2 and F8A3 are the only DE genes (Supplementary Table S7). F8A2 and F8A3 follow a similar trend in non-diseased tissues, being more highly expressed in African Americans by up to 4-fold in colon, esophagus, and thyroid (Supplementary Table S17-S18). Distribution of F8A2 and F8A3 expression is bimodal in European Americans for most cancers. Thus, part of the difference in levels of F8A2 and F8A3 expression between the two populations is due to their distribution, with low levels of expression in a subset of the European American population.

**Figure 5.**
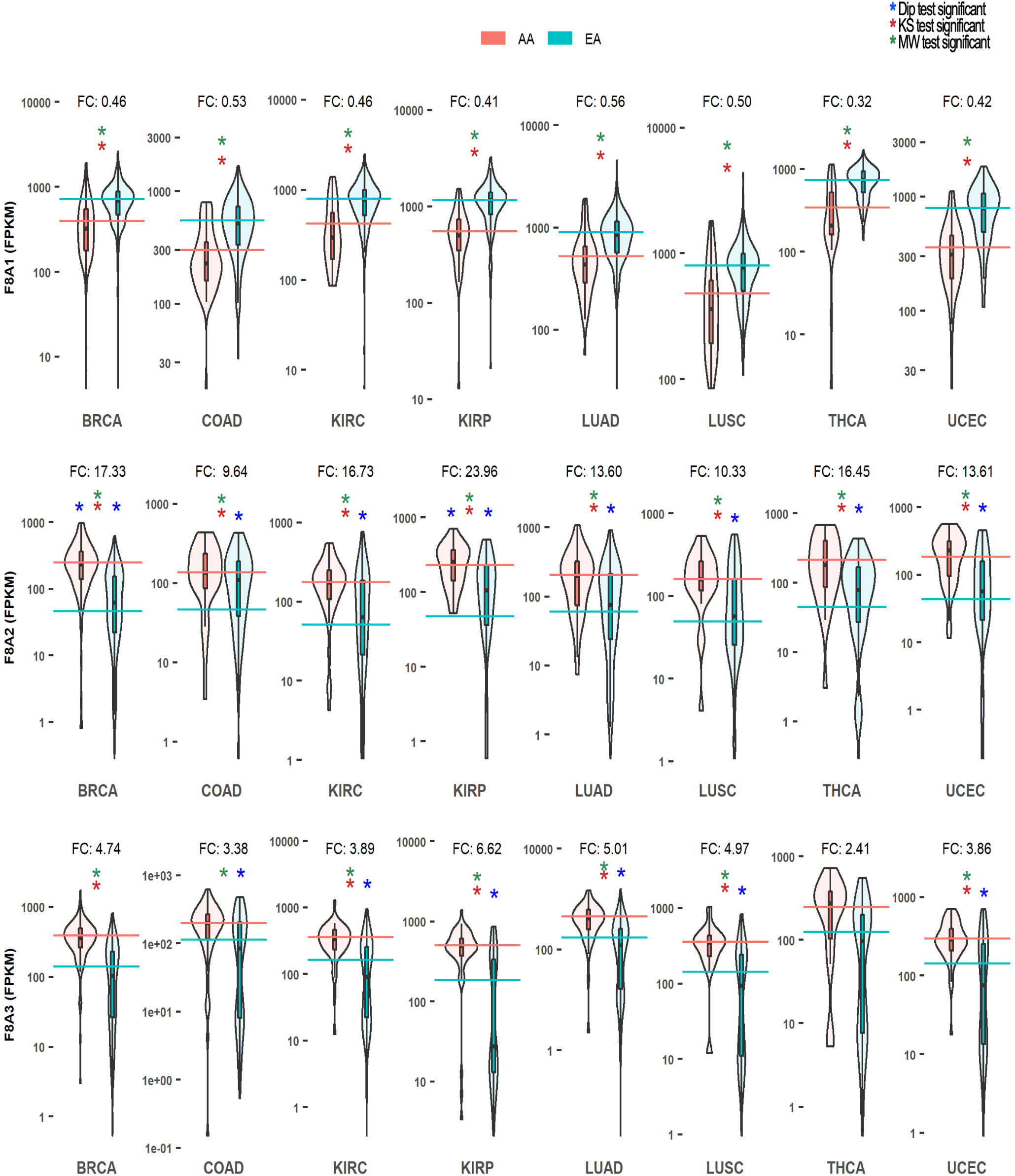
Expression of the HAP40 putative early endosome trafficking genes: **F8A1, F8A2 and F8A3** in African Americans and European Americans across eight cancer conditions. Although the function of HAP40 has not been investigated in normal individuals, this protein is a key component of Huntington’s Disease; in Huntington’s, HAP40 shifts endosomal trafficking from the microtubules to actin^55^. Violin plots summarize the expression over each tumor sample in the two populations. AA, African American; EA, European American. Horizontal lines represent mean log expression. *, Hartigans’ dip test significant (p-value < 0.05); *, KS test significant (p-value < 0.05); *, Mann-–Whitney U test significant (BH corrected p-value < 0.05).

Because of the paucity of literature on HAP40^58^ and because, to our knowledge, the relationships among F8A1, F8A2, and F8A3 genes have not been described, we investigated further the sequences, sequence variants, and the expression patterns of these genes.

The sequences of the HAP40 proteins of F8A1, F8A2, and F8A3 are identical to each other in human reference genome GRCh38.p13 (https://www.ncbi.nlm.nih.gov/assembly/GCF_000001405.39). Allele variants of HAP40 proteins encoded by F8A1, F8A2, and F8A3 were mined from The Genome Aggregation Database (gnomAD)^59^, a open database, which contains sequences of over 140,000 exomes and genomes from individuals of diverse populations as categorized by clustering of genetic features (rather than being self-reported). Individuals are assigned to one of the five major populations and to sub-populations within these. The search of gnomAD identified no variants in the HAP40s encoded by F8A2 or F8A3, and a single very rare variant (<1/1000) of F8A1 found only in European (non-Finnish) populations. The variant encodes a missense mutation (https://gnomad.broadinstitute.org/gene/ENSG00000197932?dataset=gnomad_r2_1). No structural variants were identified for HAP40 of F8A1 or F8A3; F8A2 has a rare duplication of 54 aa.

To our knowledge, F8A2 and F8A3 gene expression has not been described. We analyzed coexpression of the three F8A genes in the context of the other 18,212 genes represented in the full TCGA-GTEx dataset, using the “Pearson correlation” function in MOG. Although the F8A genes are proximately located on the X chromosome, the genes are *not* highly coexpressed with each other (| Pearson Corr. | < 0.46). Furthermore, no F8A gene is coexpressed with *any* other of the 18,212 genes represented in the dataset (| Pearson Corr. | < 0.46) (Supplementary Table S30). Of the 18,212 genes, the expression of F8A1 is most *negatively* (anti-) correlated with those of F8A2 and F8A3, with Pearson Correlations of −0.45 and −0.24, respectively (Supplementary Table S30). Mutual information (MI) can detect linear as well as complex non-linear associations, whereas Pearson’s correlation measure quantifies linear dependencies. Using the MI function in MOG, we found that F8A2 and F8A3 genes are most associated with F8A1 (Supplementary Table S30), indicating that there may be a negative interaction among these genes.

## Discussion

Genetics of human populations contribute to the propensity and severity of diseases^37, 39, 41, 41, 45, 60–65^. Sometimes the contribution is straightforward; a single allele variation found in Ashkenazi Jews, causes the vast majority of Tay-Sachs disease^66^. Sometimes it is more complex; for example, hypertension, which more prevalent in African American than European American populations^60^ in part due to detrimental APOL1 mutations that are more frequent in West African populations^62^. Despite the paucity of studies focused on Western African populations, the propensity and severity of several other diseases among this population have been attributed to genetics^41, 62, 67^.

The individual’s immune system is key to fighting viral infections. However, conversely, many COVID-19 deaths have been attributed to a cyclic over-excitement of the innate immune system. This latter process, often termed a cytokine storm, results in a massive production of cytokines and the body attacking itself, rather than specifically destroying the pathogen-containing cells^14^. Thus, people with comorbidities, the elderly, and immunosuppressed individuals, may be at a greater risk for COVID-19 morbidity and mortality because they may not respond to infection with sufficient immune response^50^ and/or because they may be more likely to develop a cytokine storm^14^. We focused on ten genes that are differentially expressed between African Americans and European Americans are implicated in these biological processes.

The most dramatic differences in gene expression were in expression of the F8A genes, F8A1 being upregulated in European Americans, and F8A2 and F8A3 being upregulated in African Americans. Each F8A gene encodes an identical HAP40 protein. F8A1, F8A2 and F8A3 genes each have a very distinct pattern of expression across the thousands of samples of tissues and cancers in the TCGA/GTEx dataset.

HAP40 function has been researched only in the context of its critical role in early endosome maturation in Huntington’s disease^56^. In Huntington’s, HAP40 forms a bridge between the huntingtin protein and the regulatory small guanosine triphosphatase, RAB5; formation of this complex reduces endosomal motility by shifting endosomal trafficking from the microtubule to the actin cytoskeleton^55^. High F8A1 expression has been reported in several conditions: Huntington’s^68^; a SNP variant for type 1 diabetes risk^69^; cytotrophoblast-enriched placental tissues from women with severe preeclampsia^70^; and mesenchymal bone marrow cells as women age^71^. Although its non-disease biology has been little explored, because of its role in early endosome motility in Huntington’s, HAP40 is considered a potential molecular target in therapy of autophagy-related disorders^72^.

Endosome motility and development play an important but complex role in the innate immune response, which can either promote or hinder the battle between SARS-CoV-2 and its human host^51, 54, 73, 74^. Coronaviruses including SARS-CoV-2 mainly enter host cells via binding to the ACE2 receptor followed by endocytosis^20, 29, 51^. Nascent early endosomes are moved along the microtubule cytoskeleton, fusing with other vesicles; varied molecules can be incorporated into the membrane or the interior^51, 54, 73, 74^. This regulated development enables diverse fates. For example, in the context of SARS-CoV-2, endosomes might release viral RNA or particles; they might merge with lysosomes and digest their viral cargo; or might fuse with autophagosomes (autophagy) and subsequently with lysosomes that digest the cargo^51, 54, 73, 74^. SARS-CoV-2 might reprogram cellular metabolism, suppressing autophagy and promoting viral replication,^75^. The cell might modify autophagy machinery to decorate viral invaders with ubiquitin for eventual destruction, activate the immune system by displaying parts of the virus, or catabolize excess pro-cytokines. Autophagy might induce cytokine signaling, which could promote protective immune response or engender a destructive storm of cytokines, inflammation and tissue damage^54^.

Cytokines and other immunomodulatory molecules, including CCL3L3, CXCL9, CXCL10, CEACAM5 and CEACAM6, were differentially expressed between African Americans and European Americans. CCL3L3 is a member of the functionally-diverse C-C motif chemokine family. It encodes CCL3, which acts as ligand for CCR1, CCR3 and CCR5 recruits and activates granulocytes; it also inhibits HIV-1-infection^76^. CCL3L3 is upregulated in younger and impoverished white males^77^. Circulating chemokines CXCL9 and CXCL10 initiate human defenses, and potentially instigate autoimmune and inflammatory diseases, by activating G protein-coupled receptor CXCR3^78–80^. CEACAM5 and CEACAM6 are members of the C-Type Lectin Domain Family. This gene family encodes a diverse group of calcium (Ca2+)-dependent carbohydrate binding proteins, several of which, including CEACAM5 and CEACAM6 have been implicated as having specific cell adhesion, pathogen-binding and immunomodulatory functions^81^. CEACAM5, a driver of breast cancer^82^ and modulator of inflammation in Crohns Disease^83^, and CEACAM6, an inhibitor of breast cancer when coexpressed with CEACAM8^84^, are both downregulated 2 to 3-fold in BRCA in African Americans.

Reactive Oxygen Species (ROS) generated in the mitochondria promote the expression of proinflammatory cytokines and chemokines, thus playing a key role in modulating innate immune responses against RNA viruses^85–89^. Mitochondrially-targeted glutathione S-transferase, GSTM1, which was more highly expressed in African Americans than European Americans, is a key enzyme in the metabolism of ROS, as well as xenobiotics including pharmaceuticals^88^. GSTM1 is induced by nuclear factor erythroid 2-related factor 2 (Nrf2), a transcription factor that integrates cellular stress signals^90–93^. Low expression of GSTM1 can lead to increased mitochondrial ROS, which may ultimately result in a cytokine storm that triggers inflammation and/or autoimmune disease. Conversely, if GSTM1 is too highly expressed, pharmaceuticals may be metabolized and thus rendered inactive, and ROS may be metabolized too rapidly to maintain a sufficient signaling role in the immune system. Allele frequencies of GSTM1 vary among Asian, African and European populations^94^; the biological significance of these alleles is being investigated^95, 96^.

CEACAM5 and CEACAM6, both downregulated 2 to 3-fold in BRCA in African Americans, are members of the C-Type Lectin Domain Family. This gene family encodes a diverse group of calcium (Ca2+)-dependent carbohydrate binding proteins; CEACAM5 and CEACAM6 have been implicated as having specific cell adhesion, pathogen-binding and immunomodulatory functions^81^. CEACAM5 is a driver of breast cancer^82^ and modulator of inflammation in Crohns Disease^83^, and CEACAM6 is an inhibitor of breast cancer when coexpressed with CEACAM8^84^.

By revealing differential expression of genes implicated in COVID-19 morbidity and mortality between African Americans and European Americans, we emphasize the importance of integrating gene expression data into the mix of factors considered in studying this pandemic. Our study indicates that, under both diseased and non-diseased conditions, many genes involved in infection, inflammation, or immunity are differentially expressed between African Americans and European Americans. One contributing explanation of the finding that disease-related genes are overrepresented among DE genes is that the selection pressure due to disease is very strong on both (ancestral) regions, but these regions have very different complements of pathogens. Humans living in Europe and those living in Western Africa would have had to evolve the ability to resist the prevalent local pathogens.

Archived expression data has tremendous value. However, studies such as this one are hampered by several factors. For example, obtaining adequate sample sizes for statistical analysis of populations is very important but difficult or expensive to address. Ethnic bias and practical factors (such as subject availability) often result in insufficient numbers of subjects from many populations to be represented in medical studies; this lack of representation prevents the development of precise prognosis or therapy based on genetics^39, 97^. Similarly, diverse socioeconomic contexts may not be well represented among the individuals sampled. Yet these are clearly a factor in disease^98^. The statistical predictor provided for the U.S. by the Robert Wood Johnson Foundation (https://www.rwjf.org/en/library/interactives/whereyouliveaffectshowlongyoulive.html) reflects the concept that “your zip code can be greater than your genetic code” (although unfortunately there is no accompanying genetic information).

Another factor that would greatly increase the utility of archived expression data is improving and extending the metadata for all (future) studies; this would be relatively simple to implement. Well-constructed metadata is key to the usefulness of data. Among the vast body of human RNA-Seq data being deposited, fields for age and gender are typically represented and available in the metadata. However, except in specialized studies, metadata on the race and ethnic heritage of the sampled individuals are often not included, or are very difficult to access. The same is even more true of fields that would provide socio-economic information, such as postal code or risk factors such as occupation. Because of the absence of socioeconomic metadata, even in GTEx and TCGA, the arguably most comprehensive RNA-Seq datasets to date, we were unable to distinguish genetic effects from environmental causes of the differences in gene expression. Without routine inclusion of diverse metadata for human ‘omics samples, data re-mining is hampered, and important information is lost.

## Conclusion

Multiple genes implicated in COVID-19 are differentially expressed in African American and European American populations. The differential expression is evident despite the fact that race is self-reported in and metadata, and that many Americans are racially admixed^41^.

Gene expression represents the interaction of genetic and environmental factors. Routine inclusion of information on ethnicity, race, postal code (as a proxy for socioeconomic condition), and profession in the metadata for each individual sampled would empower large-scale data-driven approaches to dissect the relationships between race, socio-economic factors, and disease.

By highlighting the wide-ranging differences in expression of several disease-related genes across populations, we emphasize the importance of harvesting this information for medicine. Such research will establish prognostic signatures with vast implications for precision treatment of diseases such as COVID-19.

## Methods

The MOG tool was used to interactively explore, visualize and perform differential expression and correlation analysis of genes. We downloaded the precompiled MOG project http://metnetweb.gdcb.iastate.edu/MetNet_MetaOmGraph.htm^42^, created using the data processed by Wang et. al., in which expression values have been normalized and batch corrected to enable comparison across samples^43^. This *MOG_HumanCancerRNASeqProject* contains expression values for 18,212 genes, 30 fields of metadata detailing each gene, across 7,142 samples representing 14 different cancer types and associated non-tumor tissues (TCGA and GTEX samples) integrated with 23 fields of metadata describing each study and sample.

Since the data was normalized and batch corrected, we used Mann-Whitney U test, a non-parametric test, to identify differentially expressed genes between two groups. An R script was written to perform KS and dip tests, and create the violin plots and executed via MOG interactively.

Pearson correlation values were computed, after data was *log*2 transformed within MOG, in MOG’s statistical analysis module.

Information on how to reproduce the results are available at https://github.com/urmi-21/COVID-DEA.

## Data availability

We subscribe to FAIR data and software practices^99^. MOG is free and open source software published under the MIT License. MOG software, user guide, and the *MOG_HumanCancerRNASeqProject* project datasets and metadata described in this article are freely downloadable from http://metnetweb.gdcb.iastate.edu/MetNet_MetaOmGraph.htm. MOG’s source code is available at https://github.com/urmi-21/MetaOmGraph/. Additional files are available at https://github.com/urmi-21/COVID-DEA.

## Supplementary data

Supplementary data are available at bioRxiv.

## Acknowledgements

We thank Mashette Syrkin-Nikolau, Diane Bassham, and Judy Syrkin-Nikolau for valuable discussion and comments on the manuscript. This work used the Extreme Science and Engineering Discovery Environment (XSEDE), which is supported by National Science Foundation grant number ACI-1548562. In particular it used the Bridges HPC environment through allocations TG-MCB190098 and TG-MCB200123 awarded from XSEDE. It also used the Condo Cluster at Iowa State University.

